# Acoel single-cell atlas reveals expression dynamics and heterogeneity of a pluripotent stem cell population

**DOI:** 10.1101/2022.02.10.479464

**Authors:** Ryan E. Hulett, Julian O. Kimura, D. Marcela Bolaños, Yi-Jyun Luo, Lorenzo Ricci, Mansi Srivastava

**Affiliations:** Department of Organismic and Evolutionary Biology, Museum of Comparative Zoology, Harvard University, Cambridge, MA 02138

## Abstract

Pluripotent adult stem cell populations underlie whole-body regeneration in many distantly related animal lineages. These collectively pluripotent populations of cells share some features across species, such as the expression of *piwi* and other germline-related genes. Studies of how these cells operate during regeneration are needed in diverse systems to determine how underlying cellular and molecular mechanisms of renewal and differentiation compare. Here, we sought to characterize stem cells and their dynamics in the acoel *Hofstenia miamia*, a highly regenerative marine worm with a *piwi*-expressing stem cell population called neoblasts. Transcriptome profiling at single cell resolution revealed cell types shared across postembryonic stages, including stem cells and differentiated cell types such as neural, epidermal, muscle, and digestive cells. Reconstruction of single-cell differentiation trajectories followed by functional studies confirmed that neoblasts are the source of differentiated cells and identified transcription factors needed for the formation of major cell types. Next, analysis of single-cell transcriptomes from regenerating worms showed that both differentiated cells and stem cells dynamically alter gene expression in response to amputation. Further analysis of the stem cells recovered subpopulations of neoblasts, each with specific transcriptional profiles suggesting that the majority of neoblasts are specialized to differentiated lineages, reflecting putatively lineage-primed progenitors. Notably, neoblast subsets in *Hofstenia* were identifiable consistently across postembryonic stages and also displayed differential expression dynamics in response to wounding. Altogether, these data suggest that whole-body regeneration is accomplished by the coordination of cells with distinct and dynamic transcriptomic profiles through time. Furthermore, the data generated here will enable the study of how this coordination is achieved, enhancing our understanding of pluripotent stem cells and their evolution across metazoans.

## Introduction

Regeneration is a fundamental biological process that, in most animal species, requires precise control of stem cell self-renewal and differentiation in the adult animal body. The phenomenon is widely observed across animal phyla, albeit with different animals displaying varying capacities to replace tissue (Bely & Nyberg, 2010). In vertebrates, most post-embryonic stem cells are lineage-restricted, corresponding to limited regeneration capacity in the adult body. In contrast, many invertebrates such as cnidarians, acoels, planarians, and tunicates can regrow their whole bodies after injury, and, as adults, carry a large pool of effectively pluripotent stem cells that enable regeneration of virtually any missing cell type (Lai & Aboobaker, 2018). Whether a conserved mechanism drives regulation of these stem cells across metazoans remains unclear. Thus, identification of mechanisms for the regulation of adult stem cells is central to a comprehensive understanding of whole-body regeneration.

Pluripotent stem cell populations identified in diverse invertebrates express homologs of the Piwi gene family and, in some cases, other well-known germline genes (Juliano et al., 2010; van Wolfswinkel, 2014). The planarian *Schmidtea mediterranea* can regenerate any missing tissue and has a large population of Piwi-expressing stem cells called “neoblasts” that are required for regeneration (Newmark & Sanchez Alvarado, 2000; Reddien et al., 2005). Single neoblasts transplanted into irradiated animals expand clonally (thus referred to as clonogenic-or c-neoblasts), differentiate into all tissue types of the adult animal, and restore the regenerative capacity of their hosts (Wagner et al., 2011). However, neoblasts in adult worms are heterogeneous - a large proportion are likely lineage-primed progenitors (Adler & Sanchez Alvarado, 2015; Adler et al., 2014; Cowles et al., 2013; Fincher et al., 2018; Lapan & Reddien, 2011; Plass et al., 2018; Scimone et al., 2014, 2011) and it is unknown if any one subset of the total neoblast population is clonogenic. Recent studies suggest that lineage-primed neoblasts may give rise to progeny with a different fate, i.e. seemingly lineage-restricted cells could be pluripotent (Raz et al., 2021). The Piwi-expressing stem cells of cnidarians, called i-cells, show slightly different potency in different species (Plickert et al., 2012). Within cnidarian systems, there is heterogeneity within the population of i-cells found in *Hydractinia* and *Hydra* (Plickert et al., 2012) and these stem cell populations are considered to be multipotent or effectively pluripotent (Gahan et al., 2016; Juliano et al., 2014; Rebscher et al., 2008; Siebert et al., 2019). Acoel worms are another example of an animal group that regenerate through the use of stem cells called neoblasts. Acoels hold a unique phylogenetic position that is sister to other bilaterians (Cannon et al., 2016; Hejnol et al., 2009; Mwinyi et al., 2010; I. Ruiz-Trillo et al., 2002, 1999; Iñaki Ruiz-Trillo et al., 2004; Sempere et al., 2007; Telford et al., 2003) or ambulacrarians (Kapli et al., 2021; Kapli & Telford, 2020; H. Philippe et al., 2011, 2007; Hervé Philippe et al., 2019) that can inform stem cell differentiation dynamics and provide a comparative perspective to reveal the evolution of bilaterian cell-type programs.

Here we applied single-cell RNA-sequencing (scRNA-seq) to profile cell states during postembryonic development and whole-body regeneration in the acoel *Hofstenia miamia. Hofstenia* regenerate robustly and are amenable to functional studies of regeneration. Despite the large evolutionary distance between *Hofstenia* and planarians, *Hofstenia* also have stem cells called neoblasts that are the only proliferating cell types, are required for regeneration, and express homologs of *piwi* (Gehrke & Srivastava, 2016; Srivastava et al., 2014). Constructing a *Hofstenia* single-cell atlas enabled us to identify major cell types and the transcription factors required for their differentiation from stem cells. scRNA-seq of regenerating worms revealed that stem cells and differentiated cell types respond to amputation with dynamic but cell-type specific changes in gene expression. Further, we found that the stem cell population is composed of transcriptionally distinct subtypes that respond differently to amputation.

## Results

### A single-cell transcriptome atlas of juvenile and adult worms

*Hofstenia* are maintained in the laboratory as a sexually reproducing population of hermaphroditic adults. Zygotes undergo embryonic development and hatch out as juvenile worms in about 8-9 days, with post-embryonic development into sexually mature adult worms occurring over 6 weeks (Kimura et al., 2021) (Fig. 1a). To facilitate a thorough characterization of cell types and their dynamics, we applied single-cell RNA-sequencing (scRNA-seq) using the InDrops platform (Klein et al., 2015; Zilionis et al., 2017) in whole worms at four distinct stages of post-embryonic development: 1) hatchling juveniles, worms that have newly emerged from the egg shell upon completion of embryonic development (∼8 days post lay, dpl), 2) late juveniles, worms with clear anterior and posterior bands of cream-colored pigmentation on their dorsal sides (typically 14 dpl), 3) early adults, worms with visible oocytes on their ventral sides (typically 30-40 dpl), and 4) late adults, worms that are sexually mature and have begun to produce embryos (typically >90dpl).

**Figure 1.**
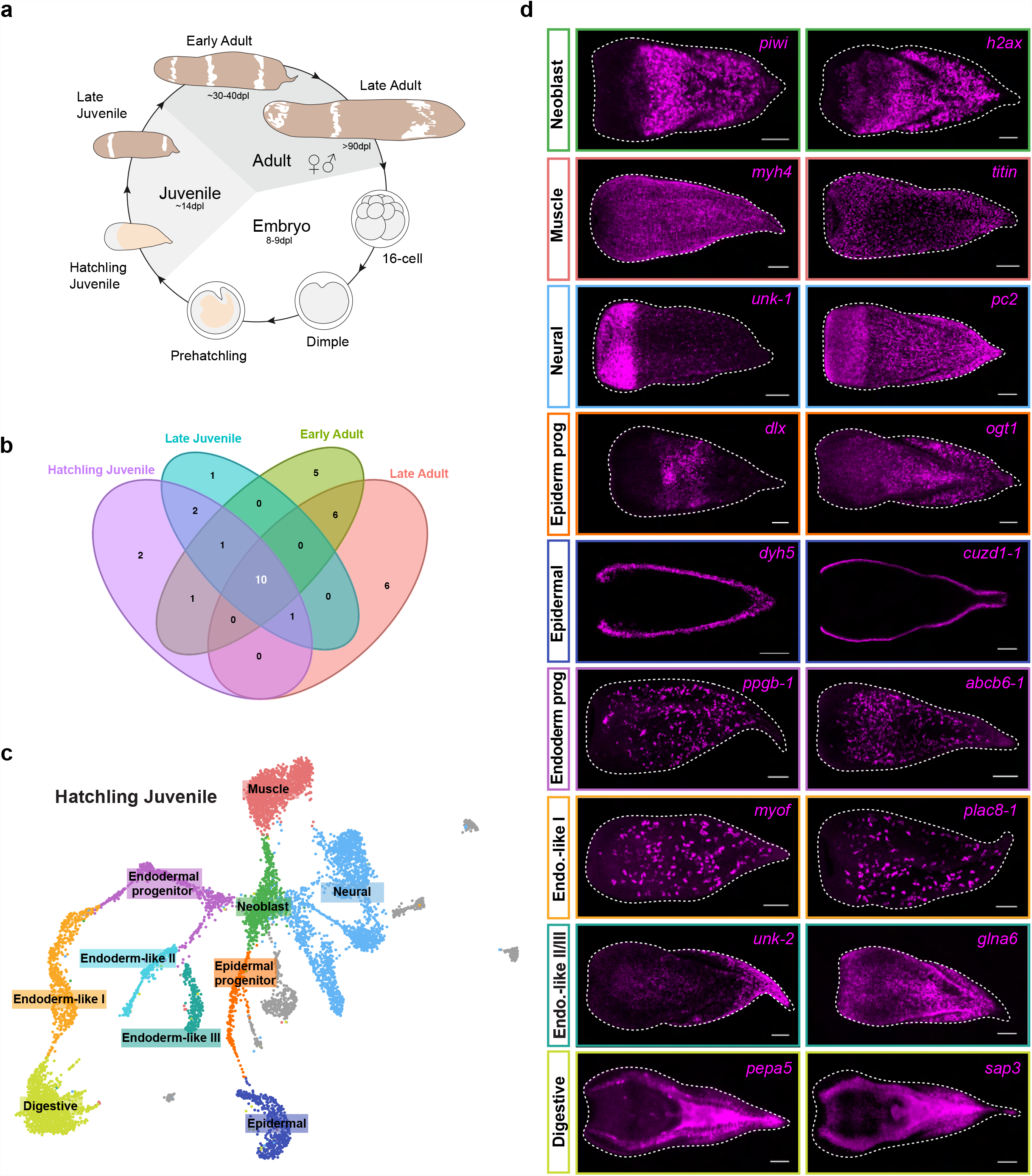
scRNA-seq reveals major cell types present across postembryonic stages. (A) Life cycle schematic of *Hofstenia miamia. Hofstenia* embryogenesis takes about 8-9 days, and postembryonic development until sexual maturity takes about 6 weeks. The postembryonic stages are divided into 4 distinct stages: hatchling juvenile, late juvenile, early adult, and late adult. (B) Venn diagram of *Hofstenia* cell types detected across different postembryonic stages. There are 10 cell types that are consistent across postembryonic development, and these were the cell types that were studied. (C) Uniform Manifold Approximation and Projection (UMAP) representation of the hatchling juvenile scRNA-seq dataset with the 10 cell types that were consistent across stages labeled. Gray cells represent cell types that were not consistent across stages. (D) *Fluorescent in situ* hybridization of selected markers corresponding to each of the identified cell types. Scale bars 100 µm.

Unsupervised clustering of data from each stage revealed variable numbers of detectable putative cell types across post-embryonic development (Fig. S1a, Table S1). To determine the identities of putative cell types, we focused on clusters that were consistently detected across all stages. We calculated Pearson’s correlation coefficients based on the transcriptional profiles for all cell clusters identified in the four postembryonic stages and conducted hierarchical clustering (Fig. S1b). We found distinct clades where cell clusters from different stages grouped together, suggesting the presence of cell types with similar transcriptional profiles across postembryonic stages. Based on this analysis and the expression/projection of marker genes, we found 10 transcriptionally distinct cell types that were consistently present during postembryonic development (Fig. 1b, Fig. S1c) and we next sought to determine the identities of these cell clusters shared across stages. We focused on these putatively shared cell types rather than stage specific cell types in order to make robust, stage-independent inferences regarding differentiation trajectories. Using the hatchling juvenile as our proxy (Fig. 1c), we identified highly expressed genes, performed gene ontology (GO) enrichment analysis (Fig. S1d), and characterized mRNA expression by fluorescent *in situ* hybridization (FISH) for each cluster shared across stages (Fig. 1d, Fig. S1e). For a given cell cluster, marker genes exhibited similar gene expression patterns, suggesting the cell clusters correspond to cell types.

We found 5 clusters corresponding to known tissue types we would expect to find in the animal by considering known cell type expression patterns in *Hofstenia*: neoblast, muscle, neural, epidermal, and digestive cells (Fig.1c-d, Fig. S1b, Fig. S1c). These 5 cell types and their associated biological functions correspond to cell types identified in the single-cell sequencing data of another acoel species, *I. pulchra*, suggesting these cell types are shared across acoels. Genes enriched in the neoblast cluster lacked expression anteriorly and labeled cells with an even distribution from the midsection along the anterior-posterior axis, to the posterior-most region of the animal. This distribution is highly redolent of the pattern observed for the known neoblast marker *piwi-1* (Srivastava et al., 2014). Neural markers were highly expressed in cells in the anterior as well as in subepidermal cells in the entire body, matching the patterns of known neural markers (Hulett et al., 2020). Epidermal marker gene expression was detectable in cells on the dorsal and ventral surfaces of hatchlings. The musculature in *Hofstenia* is well-characterized (Hulett et al., 2020; Ricci & Srivastava, 2021) and we could directly observe muscle fibers in gene expression studies of muscle-related cell cluster markers. Similarly, the markers of digestive cells were expressed in cells in the interior of the animal, resembling the pattern observed in previously-studied gut markers (Kimura et al., 2021). In further corroboration of these experimental validations of cell type identities, we also found that previously identified neoblast, neural, epidermal, muscle, and gut genes were also expressed in the corresponding cell clusters in our scRNA-seq data, supporting our cell type classifications (Table S1). Additionally, GO terms enriched in these cell clusters were also consistent with these identities, *e*.*g*., muscle system process in muscle cells, synaptic vesicle transport/assembly in neurons, cilium organization in the epidermal cell (which are ciliated in *Hofstenia*), and lipid metabolic process in gut cells (Fig. S1d).

Among the remaining cell clusters, we hypothesized that two cell clusters corresponded to lineage-primed progenitor populations related to epidermal and endodermal populations because they showed expression of both differentiated cell type markers of these lineages and of neoblast markers (Fig. S1f). Given that bona fide lineage tracing is yet to be performed in *Hofstenia*, our use of the term ‘progenitor’ will be referring to putative lineage-primed progenitor populations, which are also referred to as specialized neoblasts in planarians (Scimone et al., 2014). The markers of one of the remaining clusters were expressed in large mesenchymal cells with a scattered distribution throughout the body. Although this cell cluster was distinct from digestive cells, we found that these mesenchymal cells showed overlap in gene expression with digestive cell clusters (Fig. S1f). Thus, we hypothesize that this could be an endodermal cell type, related to gut tissue, and therefore we named it Endoderm-like I. The two remaining clusters also shared gene expression with digestive and Endoderm-like I cells, and hence we refer to them as Endoderm-like II and III, respectively.

### Identification of putative neoblast differentiation trajectories

Given that neoblasts are needed for the formation of new tissue in *Hofstenia* (Srivastava et al., 2014), we next sought to identify the molecular signatures of this process in our scRNA-seq data. We found the topology of the Uniform Manifold Approximation and Projections (UMAPs) to be very consistent across the four postembryonic stages, always showing a branching structure radiating from a center. Specifically, neoblasts were located at the center, lineage-primed progenitor cells (namely the putative endodermal and epidermal) at the branch points, and mature cell types at the tips of the branches (Fig. 1c, Fig. S1a, Fig. S1c, Fig. S1d). We also found that while the clustering parameters did not recover distinct clusters for muscle and neural specialized neoblasts, *i*.*e*., cells with expression of both neoblast and differentiated cell markers, *piwi-1*^+^ cells with muscle and neural gene expression were present in our dataset, often observable at the boundary between the neoblast cluster and the muscle/neural clusters. Although dimensionality reduction approaches such as UMAP do not show transcriptional trajectories of cells (Kharchenko, 2021), our observations of the UMAP topology is highly suggestive of putative differentiation trajectories we would expect to see in *Hofstenia*. However, explicit trajectory inference tools are required to test this hypothesis. Thus, we sought to assess whether genetic effectors of cellular differentiation could be identified using a trajectory inference tool called URD (Farrell et al., 2018).

To uncover trajectories, we focused on the hatchling juvenile stage dataset. We set the neoblast cluster as the root, and for tips we selected clusters that are molecularly consistent with terminally differentiated cell types, *i*.*e*., they did not show enrichment of neoblast-associated genes or GO processes. The placement of cells along the branches of this tree can be treated as a hypothesis of differentiation trajectories. We considered the genes enriched in cells at the branch points of the URD tree to identify transcription factors (TFs) that could be regulating these differentiation trajectories. We assessed genes based on several criteria:1) they encoded TFs, 2) they were significantly enriched in branch points, and 3) their homologs had known functions in differentiation of tissues in other research models. Using this approach, we identified eight TFs as candidate regulators for muscle *(foxF* and *six1)*, neural *(vax* and *nkx2-1)*, epidermal *(foxJ1* and *dlx)*, and digestive/endodermal (*foxA, ikzf-1)* cell differentiation (Fig. 2a, Fig. S2a, Fig. S2g, Table S2). Projection of these candidate TFs onto the UMAP revealed that their transcripts were expressed either in the differentiated cell clusters and/or in specialized neoblasts (Fig. 2a, Fig. S2a). The two cell clusters we had hypothesized to be putative lineage-primed progenitors for the epidermal and endodermal lineages were placed in close relationship to differentiated epidermal and digestive cells, respectively, corroborating our hypothesis from clustering of the scRNA-seq data (Fig. S2b).

**Figure 2.**
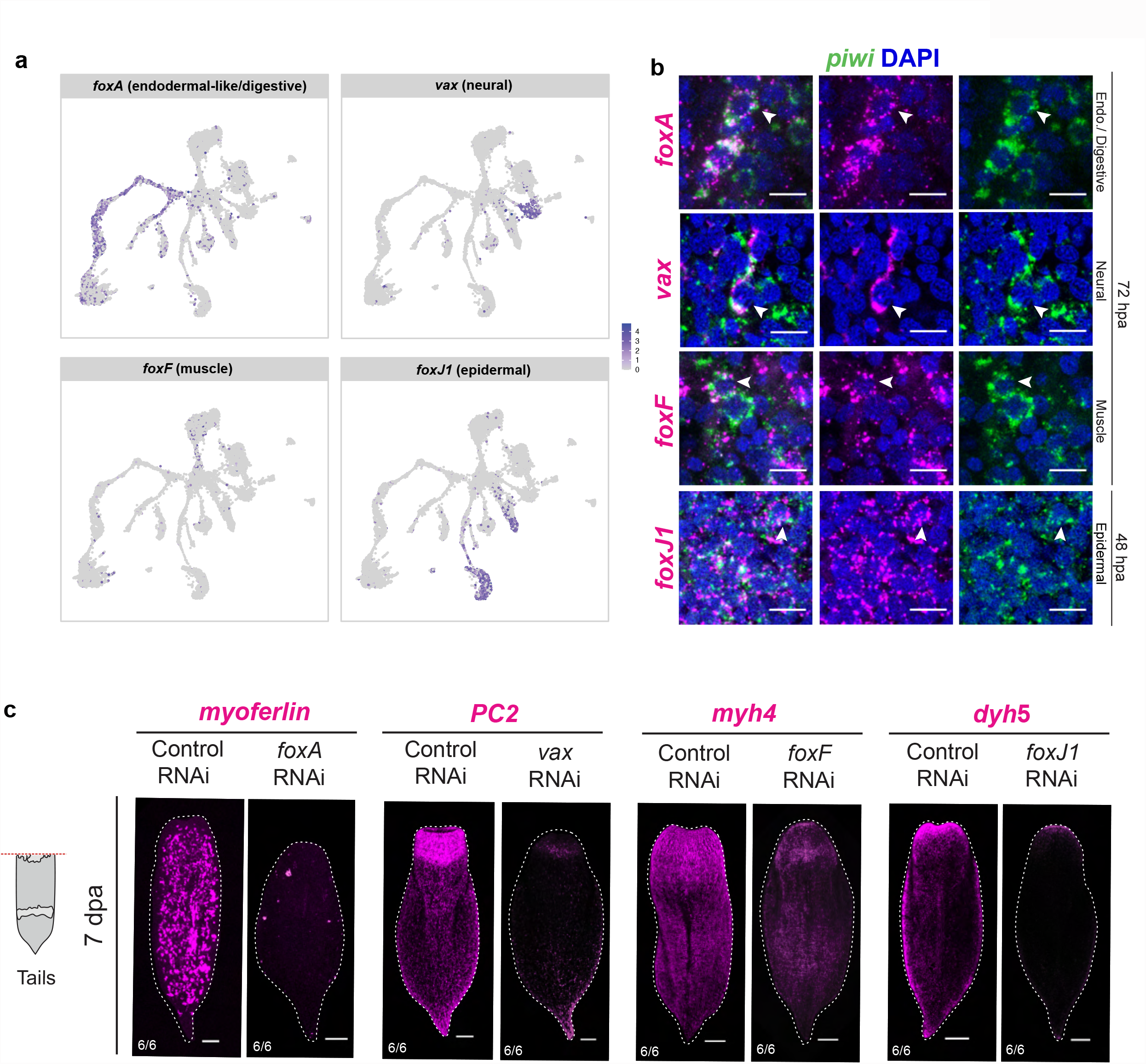
Transcription factors expressed in putative specialized neoblast populations are required for the differentiation of major cell types. (A) UMAP projection of transcription factors expressed in putative differentiation trajectories from URD analysis. All transcription factors identified were found to be expressed in their corresponding differentiated cell types and/or putative specialized neoblast populations. (B) Double FISH of the candidate transcription factors with the stem cell marker *piwi-1* during regeneration for putative endodermal-like/digestive, neural, muscle, and epidermal specialized neoblasts. Co-expression of *piwi-1* and candidate transcription factors (denoted by white arrowheads) was detected at 72 hours post amputation (hpa) for putative endodermal/digestive, neural, and muscle specialized neoblasts while putative epidermal specialized neoblasts were detected at 48 hpa. Scale bars 10 µm. (C) RNAi of putative specialized neoblast transcription factor leads to loss of differentiated cell type expression during regeneration. Regenerating tail fragments are shown 7 day post amputation (7dpa) and corresponding head fragments are in Supplementary Figure 2d. Scale bars 100 µm.

We hypothesized that if these TFs are expressed in putative specialized neoblasts that are in transition to acquiring a terminally differentiated identity, 1) their expression should be detectable in *piwi-1*^+^ cells (neoblasts), and 2) disrupting their gene expression should result in the elimination of corresponding differentiated tissues. We first sought to determine if these candidate genes are co-expressed with the pan-neoblast marker *piwi-1* in regenerating worms, as amputation forces the production and differentiation of new tissue. We found that *piwi-1*^+^/TF^+^ cells were detectable starting at 72 hours post amputation (hpa) for most TFs, and at 48 hpa for *foxJ1* (Fig. 2b and Fig. S2c) for the epidermal lineage. We were also able to detect most *piwi-1*^+^/TF^+^ cells, albeit at lower numbers, in intact animals (Fig. S2c), suggesting that differentiation pathways in homeostasis mirror those in regeneration. We next conducted functional studies using RNA interference (RNAi) to determine whether these TFs were regulators of differentiation. We assessed this process during regeneration because amputating the animal forced cells to undergo differentiation to replace missing cell types and tissues. In seven out of eight gene knockdowns, animals formed unpigmented outgrowths (blastemas) and regenerated heads and tails, recapitulating wild type external morphology (Fig. S2d). Notably, *foxA* RNAi animals were the only fragments that showed a visible phenotype with failure to regenerate head blastemas and pointed tails (Fig. S2d). FISH for markers of the corresponding differentiated cell types confirmed loss or reduction of endodermal/digestive (*myoferlin*), neural (*pc2*), muscle (*myh4*) and epidermal (*dyh5*) cells, respectively in *foxA, vax, foxF* and *foxJ1* RNAi animals (Fig. 2c, Fig. S2e). Additionally, the knockdowns for the other four transcription factors *(ikzf-1, nkx2-1, six1* and *dlx*) resulted in reduction of expression of the corresponding differentiated cell type markers relative to control RNAi and the knockdown of *FoxA* resulted in loss of digestive cell types (Fig. S2f). Together these data show that specific TFs that are expressed in subsets of *piwi-1*^+^ cells during regeneration and homeostasis are required for the formation of distinct differentiated tissues. Given that *piwi-1*^+^ cells are the only proliferative cells and are needed for regeneration in *Hofstenia*, this work reveals the differentiation pathways required for the formation of new tissues from neoblasts.

### Gene expression dynamics during regeneration

With single-cell transcriptional profiles for the major cell types in hand, we next asked whether different cell populations behave similarly or differently during regeneration in terms of gene expression dynamics. In order to evaluate cell-type specific responses to wounding, we generated scRNA-seq data from combined regenerating head and tail fragments at 7 timepoints (0, 6, 24, and 72 hours post amputation, hpa, and 8, 17, and 29 days post amputation, dpa). Unsupervised clustering of a dataset that integrates all time points recovered the major cell types identified in the postembryonic stages and the UMAP plot showed similar structure with neoblasts at the center and differentiated cell types at the periphery (Fig. 3a, Fig. S3a, Table S3). Each cluster within the UMAP was composed of cells from each of the regeneration time points, with no clearly identifiable regeneration-specific cluster (Fig. S3b).

**Figure 3.**
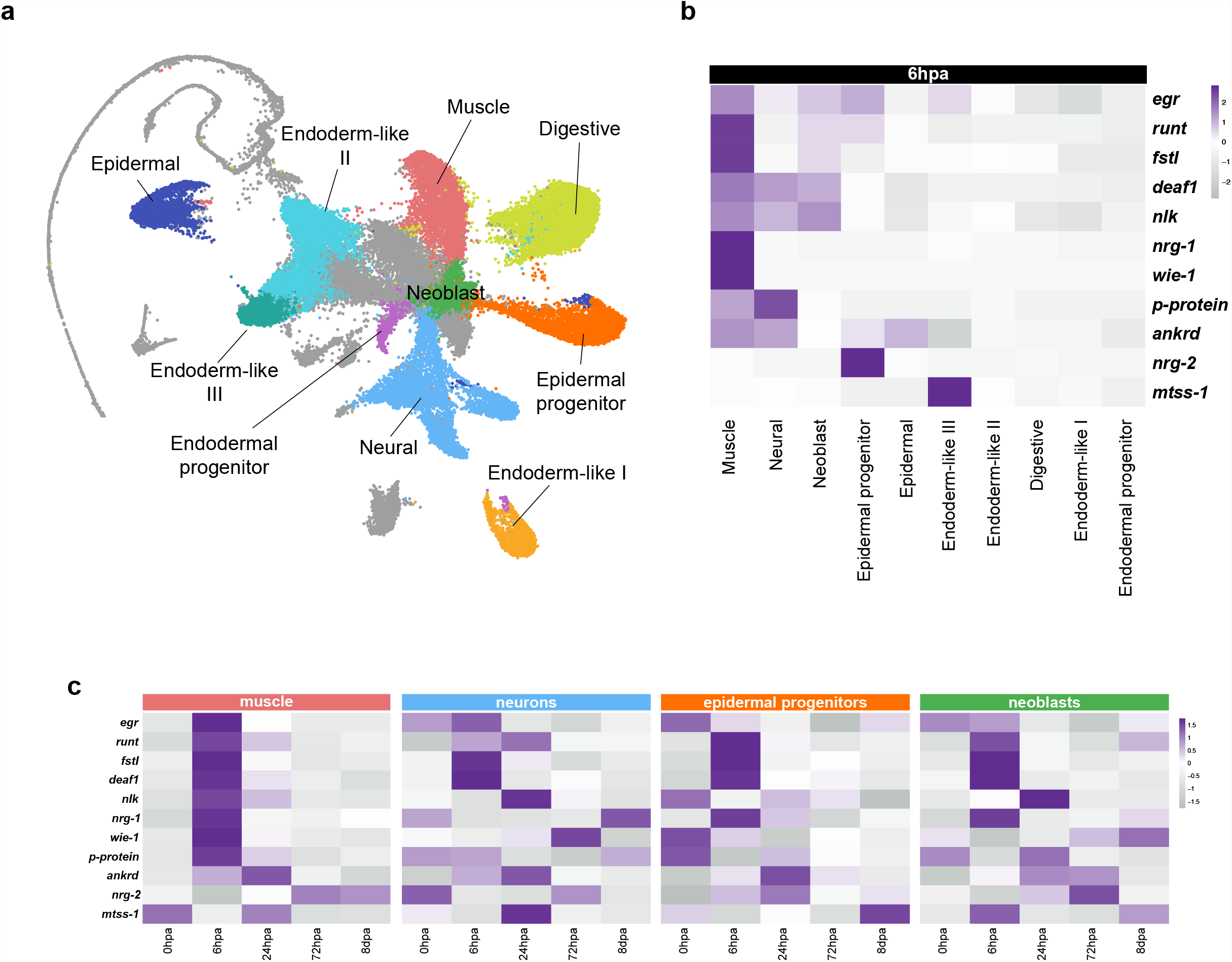
scRNA-seq during regeneration reveals cell-type specific responses during regeneration. (A) Major cell types recapitulated during regeneration. Integrated regeneration UMAP of scRNA-seq data from the following time points: 0hpa, 6hpa, 24hpa, 72hpa, 8dpa, 17dpa, and 29dpa. Ten major cell types identified during postembryonic development are recapitulated in the regeneration data, with neoblasts located in the center and differentiated cell types radiating out. (B) Cell-type specific expression of Egr-GRN at 6hpa. Cells from the integrated regeneration data were subsetted to focus on the 6hpa population and a heatmap depicting average expression was generated. All Major cell-types were populated with cells from 6hpa. We identify expression of Egr-GRN members in muscle, neural, neoblast, epidermal lineage-primed progenitors, epidermal, and endodermal-like III populations. We see either a lack of expression or down-regulation of Egr-GRN members in endodermal-like II, digestive, endodermal-like I, and endodermal lineage-primed progenitors. (C) Dynamic expression of Egr-GRN members reveals temporal cell-type specific responses during regeneration. Heatmap of the muscle population shows dynamic expression of nearly every Egr-GRN member at 6hpa. Heatmaps of the neural, epidermal lineage-primed progenitors, and neoblast populations show subsets of the Egr-GRN expressed at both 6hpa and 24hpa.

Previous work defined a gene regulatory network (GRN) controlled by the zinc finger transcription factor Egr (Egr-GRN), that is induced upon amputation in *Hofstenia* (Gehrke et al., 2019). Given that wound induction initiates the process of regeneration, which ultimately requires the coordination of multiple tissue types including neoblasts, we investigated the cellular context of this pathway. We used this GRN as a focal point to assess if the entire Egr-GRN or its components were upregulated in the many cell types in the single-cell atlas at 6hpa (Fig. 3b, Fig. S3c). Notably, the majority of genes in the Egr-GRN showed high expression in muscle cells at 6 hpa, a time point when GRN members show peak expression in bulk transcriptome data and in whole-mount gene expression studies (Gehrke et al., 2019). Strikingly, the majority of endodermal cell types, including digestive cell types, showed very little expression of any genes in the Egr-GRN. Other cell types (neural, neoblast, epidermal progenitors, epidermal cells, and endodermal-like II) expressed some Egr-GRN members. Overall, it appears that muscle, neurons, epidermal progenitors, and neoblasts respond to injury, while endodermal tissues have a relatively lower response. Interestingly, some Egr-GRN members showed cell type specificity, e.g. *nrg-1* and *wie-1* were only highly expressed in muscle. These observations suggest that Egr is not wound-induced in all cell types and that it upregulates different genes in different cell types.

To assess whether the expression of Egr-GRN members at 6 hpa was upregulated relative to 0 hpa and to identify temporal expression patterns of the GRN in specific cell types, we subset the muscle, neuron, epidermal progenitors, and neoblast populations individually from the integrated regeneration UMAP and looked at how the Egr-GRN member expression changed throughout time upon amputation to 8 dpa (Fig. 3c). All but two GRN members (*nrg-2* and *mtss-1*) showed robust upregulation from 0 to 6 hpa in muscle. Neurons, epidermal lineage-primed progenitors, and muscle showed clear upregulation of *runt, fstl, deaf1*, and *ankrd*, pointing to these genes as core members of the Egr-GRN that are upregulated in any cell that has *egr* expression upon amputation. Most core Egr-GRN genes, namely *runt, fstl*, and *deaf1* showed peak expression at 6 hpa in all cell types, whereas *ankrd* peaked at 24 hpa (except for *runt* showing peak expression in neurons at 24hpa). Some of the genes, which showed peak expression at 6 hpa in muscle, were upregulated more gradually in other cell types, e.g. *nlk* showed peak expression at 24hpa in neurons and neoblasts. This suggests that, in addition to determining which target genes get activated, the cellular context may impact the temporal dynamics of their activation.

### Heterogeneity of neoblasts

We noted that our dataset did not show a substantial upregulation of *egr* in the neoblast cluster relative to the levels observed in other cell types such as muscle (Fig. 3b, Fig. 3c, Fig. S3c), contrary to previously reported expression of *egr* mRNA in *piwi-1*^+^ cells in amputated worms (Gehrke et al., 2019). Given the well-studied heterogeneity of neoblasts in planarians (Fincher et al., 2018; van Wolfswinkel et al., 2014; Zeng et al., 2018), we asked whether heterogeneous induction of the Egr-GRN in specialized neoblast populations could underlie the lack of signal for wound-induced *egr* expression in our dataset. Therefore, we next sought to identify heterogeneity within the neoblast population.

We performed sub-clustering of the neoblast population from the integrated regeneration dataset to determine whether transcriptionally-distinct subsets of neoblasts could be identified. We recovered 11 neoblast subpopulations, each with distinct transcriptional signatures (Fig 4a). Seven of 11 clusters showed enriched expression of markers of differentiated types and/or of transcription factors we had identified as markers of specialized neoblasts. *tpm3*^+^ cells showed expression of markers of differentiated muscle cells. Based on functional studies of previously mentioned TFs, we identified 4 clusters we hypothesize represent epidermal (*dlx*^+^), muscle (*foxF*^+^), and endodermal (*ikzf-1*^+^ and *foxA*^+^) cells that likely represent lineage specialized neoblasts. Two of the remaining clusters highly expressed *sox4* and *traf2* and projections of these genes onto the hatchling juvenile UMAP showed expression in cells connecting to the central neoblast cluster and differentiated neuron clusters, indicating they may be neural specialized neoblasts (Fig. S4a). One cluster showed high expression of *boule-like* (*boll*), a gene known in other metazoan systems to be required for the specification of germ cells (Shah et al., 2010; Xu et al., 2001). *boll* is highly expressed in likely germline cells in our early and late adult datasets, and we found it to mark two populations of cells that correspond to the location of germline in FISH studies of *Hofstenia* (Fig. S1e), therefore we inferred that *boll*^*+*^ cells are possibly specialized neoblasts that are primed to make germline. One cluster showed high expression of the histone H3 variant *h3*.*3*, but no other markers that corresponded to differentiated tissues. Two clusters failed to reveal any marker genes that showed specific expression in those cells when projected back onto the UMAP, and therefore we referred to them as unknown-I and unknown-II.

**Figure 4.**
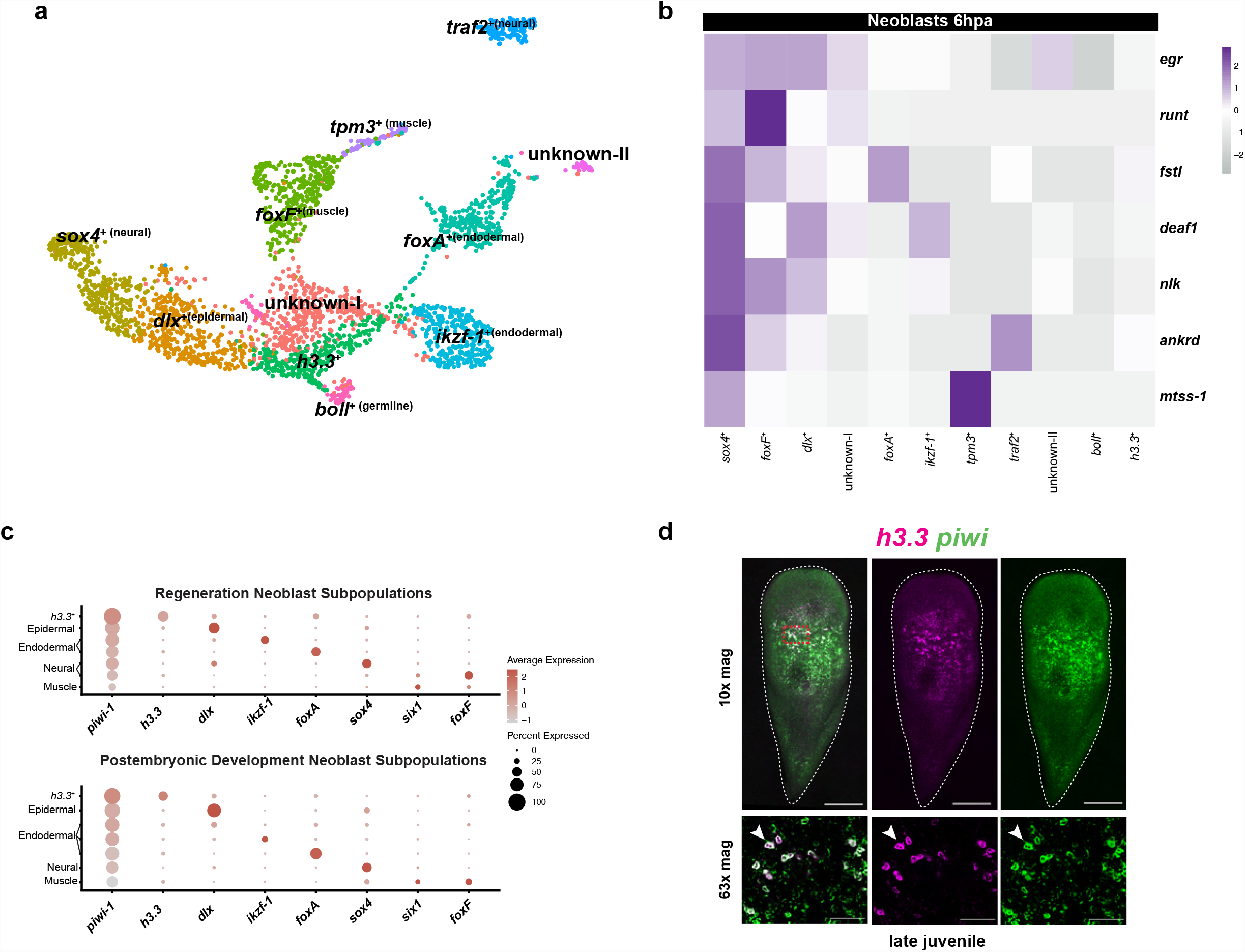
Neoblast subtype dynamics during postembryonic development and regeneration. (A) UMAP of neoblasts derived from the integrated regeneration dataset show subpopulations. Clusters are annotated with differentially expressed marker gene. (B) Heatmap of neoblast subpopulations at 6hpa showing dynamic expression of Egr-GRN members. The *sox4*^+^ neoblast subpopulation expresses every Egr-GRN member. Every other subpopulation expresses a suite of the Egr-GRN except for the *boll*^+^ which we hypothesize is a specialized germline neoblast and the *h3*.*3*^+^ subpopulation which shows downregulation of many members. (C) Dot plots of both the postembryonic development neoblast subpopulations and the regeneration neoblast subpopulations. Neoblast subpopulations that are consistently present in both the postembryonic and regeneration datasets are shown on the y-axis. Genes shown on the x-axis represent a major marker of each of these subpopulations. Strikingly, the *h3*.*3*^+^ neoblast subpopulation has high expression of *piwi-1*, relative to other specialized neoblast populations. (D) FISH reveals that *h3*.*3* (magenta) labels large mid-body cells that co-express (denoted by white arrowhead) *piwi-1* (green). 63x region denoted by red dotted square. Scale bars 200 µm for 10x magnification and 50 µm for 63x magnification.

Next, we assessed the expression of the Egr-GRN members in these neoblast subclusters and found heterogeneous expression across nearly all cell clusters at 6 hpa (Fig. 4b). Three genes of the Egr-GRN showed no or low expression in most cells across the neoblast subclusters, so they were removed from the analysis (Fig. S4b). Strikingly, one subset, the *sox4*^+^ cells (likely neural specialized neoblasts) showed robust expression of all other Egr-GRN genes (*egr, runt, fstl, deaf1, nlk, ankrd, mtss-1*). Several of these genes were also expressed in the *foxF*^+^ cells (likely muscle specialized neoblasts) and in the *dlx*^+^ cells (likely epidermal specialized neoblasts). Notably, one cluster, with *h3*.*3*^+^ cells, stood out as having very low expression of all GRN members at 6hpa.

Given that the neoblast subsets showed heterogeneous responses during regeneration, we sought to gain a deeper understanding of these subtypes. We asked whether these subsets were unique to certain time points during regeneration and whether they could be identified in intact worms. To assess whether these subclusters were found across stages, during both regeneration and development, we utilized hierarchical clustering and calculated the Pearson correlation coefficient and subclustering of the scRNA-seq data. We found neoblast subclusters associated with muscle (*foxF*^+^), neurons (*sox4*^+^), epidermis (*dlx*^+^), and endoderm (*ikzf-1*^+^, *foxA*^+^) as well as *h3*.*3*^+^ cells present in multiple stages of regeneration and postembryonic development (Fig. 4c, Fig. S4c, Fig. S4d, Fig. S4e).

Strikingly, *h3*.*3*^+^ neoblasts showed high levels of *piwi-1* expression relative to all other specialized neoblast subsets that are found across post embryonic and regeneration time points (Fig. 4c). Given that the *h3*.*3*^+^ cells are the only subset that is consistently present in all datasets from worms during postembryonic development and regeneration (Fig. S4e), remains unassigned to a lineage specialized neoblast of differentiated tissue, and was the only neoblast cluster that failed to activate any Egr-GRN gene, we sought to characterize this marker and this cell population further. FISH of *h3*.*3* revealed its expression to be almost exclusively in a population of large cells present in the midsection relative to the anterior-posterior axis of the worm. Double-FISH revealed that these cells also expressed *piwi-1*, corroborating *h3*.*3*^+^ cells as a bona fide subset of *piwi-1*^+^ cells (Fig. 4d). In addition to having enriched expression of *h3*.*3*, a histone H3 variant, these cells also express *foxO* (Fig. S4f), both genes known for their roles in the maintenance, proliferation, and cell cycle control of stem cells in other metazoan lineages (Boehm et al., 2012; Jang et al., 2015; Jullien et al., 2012; Miyamoto et al., 2007; Murdaugh et al., 2021; Sakai et al., 2009; Tothova & Gilliland, 2007; Zhang et al., 2011). These data are suggestive that *h3*.*3*^+^ cells could represent an unspecialized subset of neoblasts, but further work is needed to rule out the alternative hypotheses including these cells being a temporary cell state that many specialized neoblasts undergo.

## Discussion

The single-cell transcriptome dataset analyzed here represents an experimentally-corroborated catalog of cell types during postembryonic development and regeneration in the acoel *Hofstenia miamia*. We found that major cell types were consistently identifiable throughout these processes and included differentiated cells such as muscle, neural, epidermal, and endodermal cells as well as stem cells consisting of distinct neoblast subpopulations. We also provide functional validation of transcriptional regulators required for differentiation of neoblasts into terminally differentiated cell types. These data can provide insights into both the mechanisms of stem cell regulation and the evolution of cell type differentiation.

The transcriptome profiles for cell types and differentiation pathways in our dataset provide an important point for comparison to study the evolution of major bilaterian cell types. Muscle, neural, and digestive cells in *Hofstenia* expressed homologs of well-characterized genes that are known to have important functions in these cell types in other bilaterians. For example, muscle cells express myosins and troponin, neurons express prohormone convertases and Trp channels, and gut cells express peptidases and cathepsins. However, the expression profiles of epidermal cells in *Hofstenia* did not reveal obvious shared differentiated markers with the epidermal cells of other bilaterians found within the Nephrozoa. Previous work in the acoel *I. pulchra* identified similar cell types to those we found in *Hofstenia* based on enrichment of genes associated with biological functions (Duruz et al., 2021). In addition to uncovering these molecular functions in the corresponding *Hofstenia* cell types, our dataset recovered transcription factors associated with these cell types. We noted that these transcription factors, needed for the formation of differentiated cell types in *Hofstenia*, were homologs of known regulators of these cell types during development and/or regeneration in other bilaterians. *foxF*, a muscle regulator in *Hofstenia*, is also required for the regeneration of non-body wall muscle in planarians (Scimone et al., 2018). *Six1* homologs have well established roles in muscle and muscle progenitor differentiation (Andrikou & Arnone, 2015; Laclef et al., 2003; Niro et al., 2010; Wu et al., 2014), including during regeneration (Le Grand et al., 2012). *vax* and *nkx2* homologs, which are required for neural regeneration in *Hofstenia*, are well known neural TFs in bilaterians, and an *nkx2* homolog is needed for the maintenance of cholinergic, gabaergic, and octopaminergic neurons in the planarian central nervous system (Currie et al., 2016). *FoxJ* is a regulator of ciliated epidermal cell types across eukaryotes, and specifically results in aberrant regeneration of the epidermis in planarians (Fritzenwanker et al., 2014; Vij et al., 2012), consistent with its role in regeneration of the *Hofstenia* epidermis. *dlx*, another epidermal factor in *Hofstenia*, has homologs expressed in surface ectoderm across deuterostomes and is needed for differentiation of epidermal cells in vertebrates (Morasso et al., 1996; Panganiban & Rubenstein, 2002). *foxA* homologs are required for foregut/pharynx formation in many species (Adler et al., 2014; de-Leon, 2011; Fakhouri et al., 2010; Hejnol & Martindale, 2008; Kiefer et al., 2007; Mango et al., 1994; Updike & Mango, 2006), and we found the *foxA* homolog to be required for endodermal and digestive tissue in *Hofstenia*. Overall, differentiated cell types in *Hofstenia* appear to have clear molecular correspondence with their counterparts in bilaterians, including in terms of the underlying transcriptional regulatory pathways for the formation of these tissues.

The single-cell atlas enabled us to probe the transcriptional wound response with cell-type resolution. We found that a major wound-induced GRN that is required for regeneration is upregulated in a subset of cells, with the most robust upregulation of the majority of GRN member genes occurring in muscle. This is consistent with observations of wound-induced gene expression in planarian muscle (Cloutier et al., 2021; Petersen & Reddien, 2009; Scimone et al., 2014) and of muscle being an important regulator of regeneration (Raz et al., 2017; Scimone et al., 2017; Witchley et al., 2013). Other tissues showed upregulation of subsets of the GRN and also differed from muscle in the timing of upregulation of some genes, suggesting that the GRN has distinct dynamics and functions depending on the cellular context. We did not detect regeneration-specific clusters in our integrated regeneration data sets but we did identify the varied expression of the Egr-GRN in differentiated tissues, reminiscent of how post-mitotic cells respond to injury in the transient regeneration activated cells (TRACs) populations in *Schmidtea mediterranea* (Benham-Pyle et al., 2021). These data reiterate the importance of coordination across cell types to mediate whole-body regeneration and serve as an important resource for future work toward understanding these mechanisms.

Given the observation of *piwi*-expressing cells in many distantly-related animals, our work, together with data from planarian stem cells, suggests that we should expect *piwi*-expressing cells in other animals to be transcriptionally heterogeneous. To understand the evolution of this cell type, *piwi*-expressing stem cells should be compared not as a monolithic type, but as a mixed population that may show nuanced differences across species. Similar to planarian neoblasts, *Hofstenia* neoblasts express *piwi* homologs and are needed for regeneration (Srivastava et al., 2014). We now extend this comparison by showing how they differentiate into postmitotic cells during homeostasis and regeneration, and that *Hofstenia* neoblasts include subsets that correspond to likely specialized neoblasts. Notably, our data revealed one subset that was not assignable to a clear specialized neoblast type and showed distinct characteristics relative to other neoblast subsets. These *h3*.*3*^+^ cells are a subset of *piwi*^+^ cells that do not upregulate wound response genes and show high *piwi* expression. Future work will elucidate if this population represents a subset with distinct functions and dynamics from specialized neoblasts. In addition to revealing these stem cell regulatory mechanisms in *Hofstenia*, the data generated in this study will facilitate rigorous comparisons of stem cell dynamics across species.

**Supplementary Figure 1.**
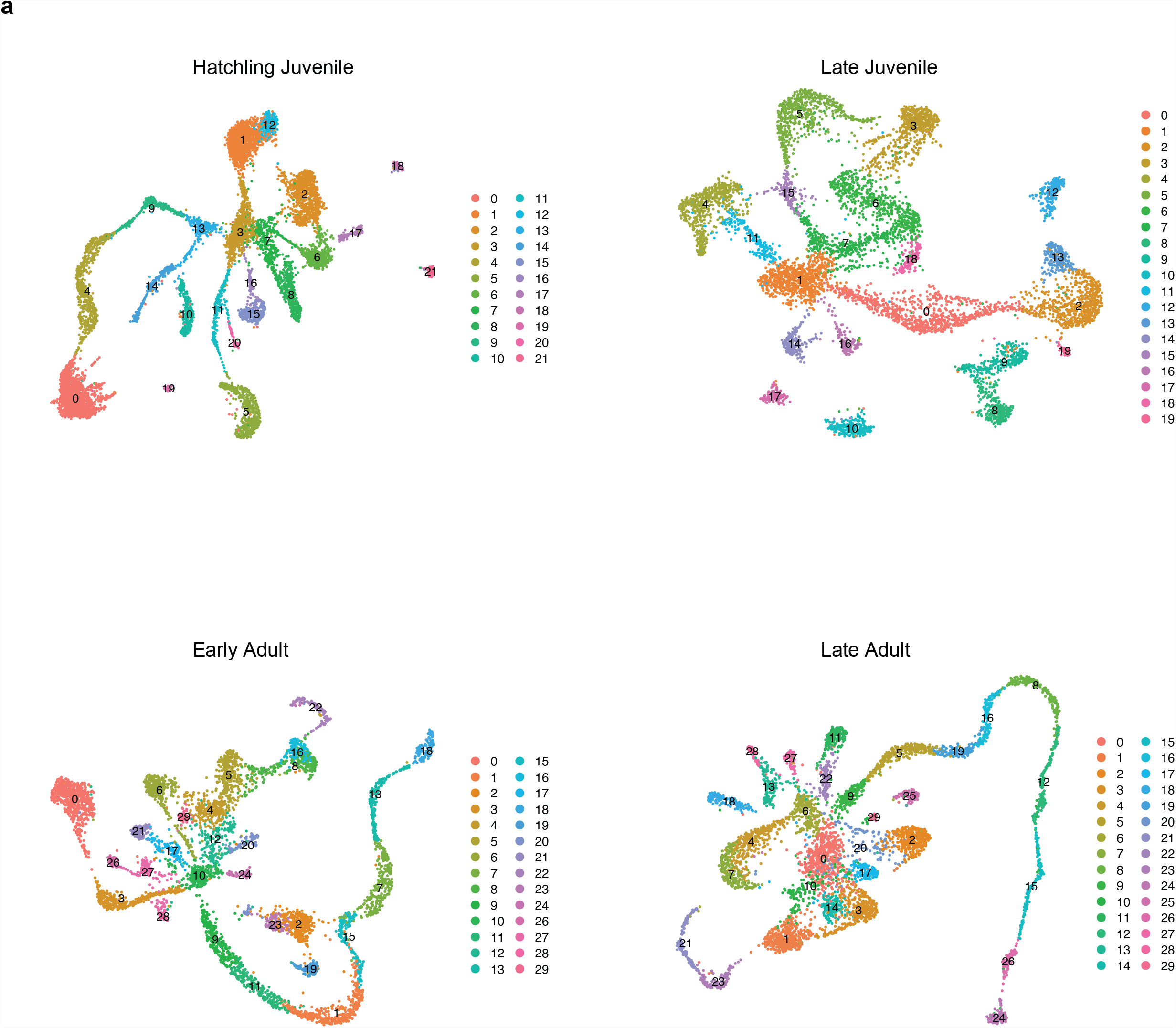
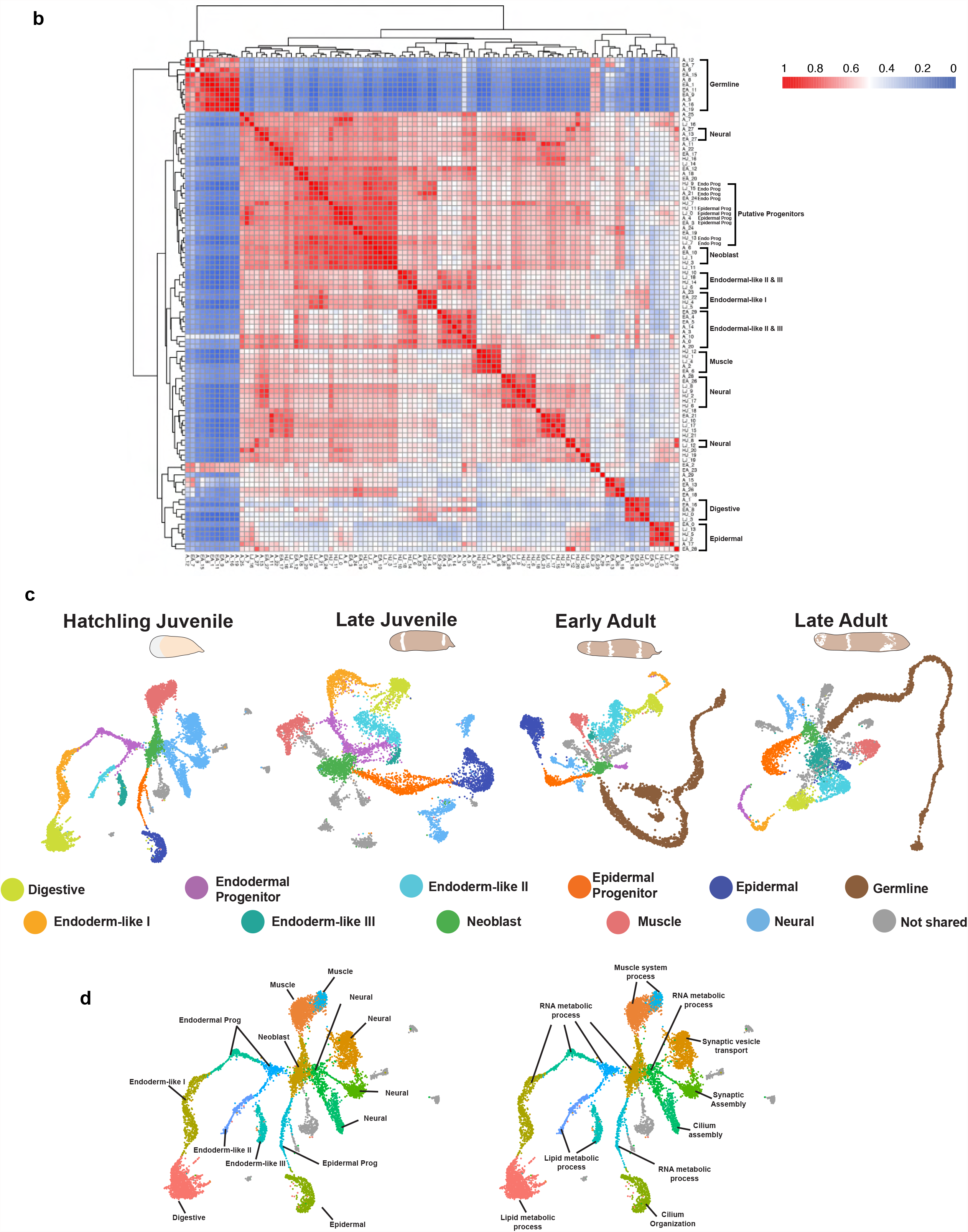
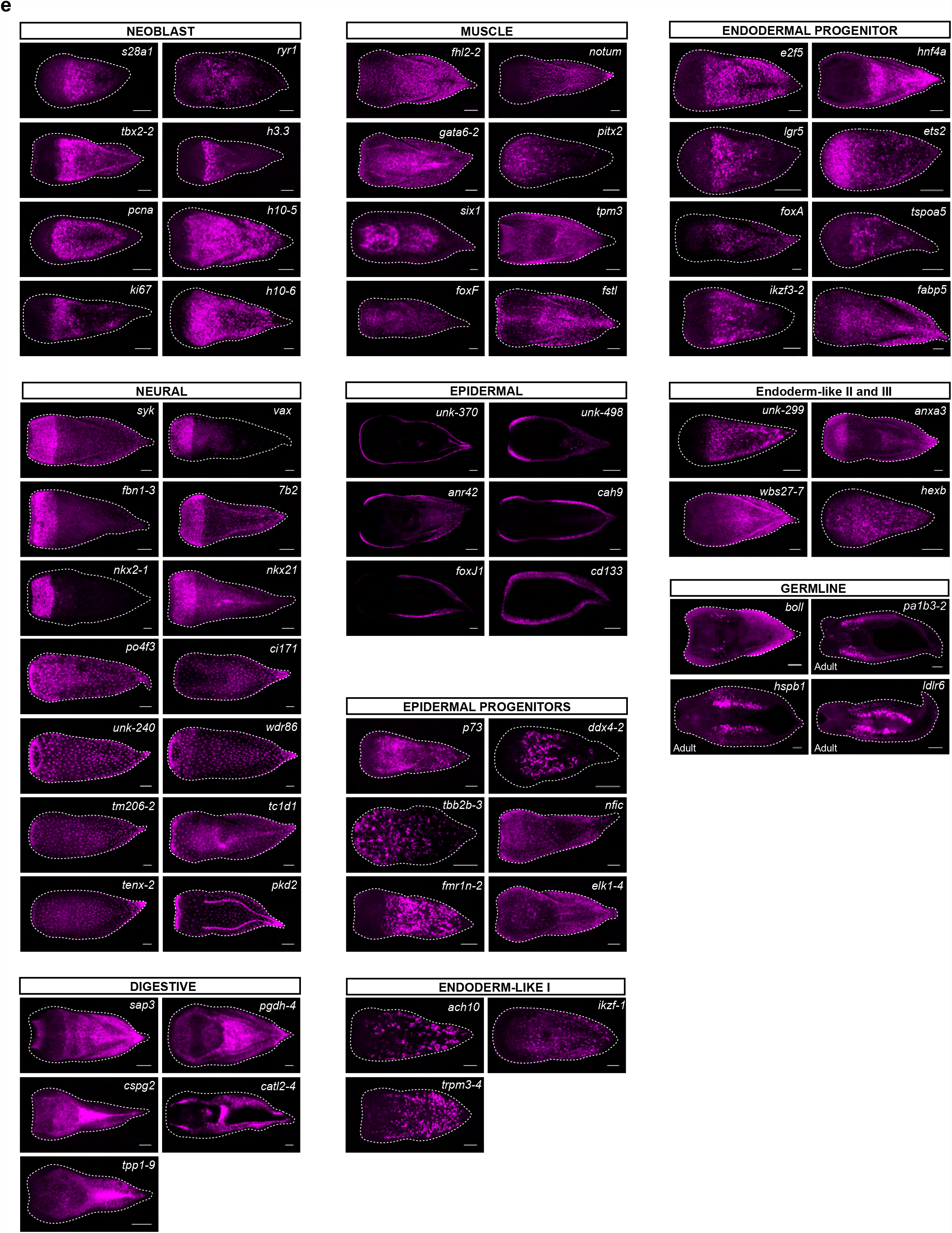
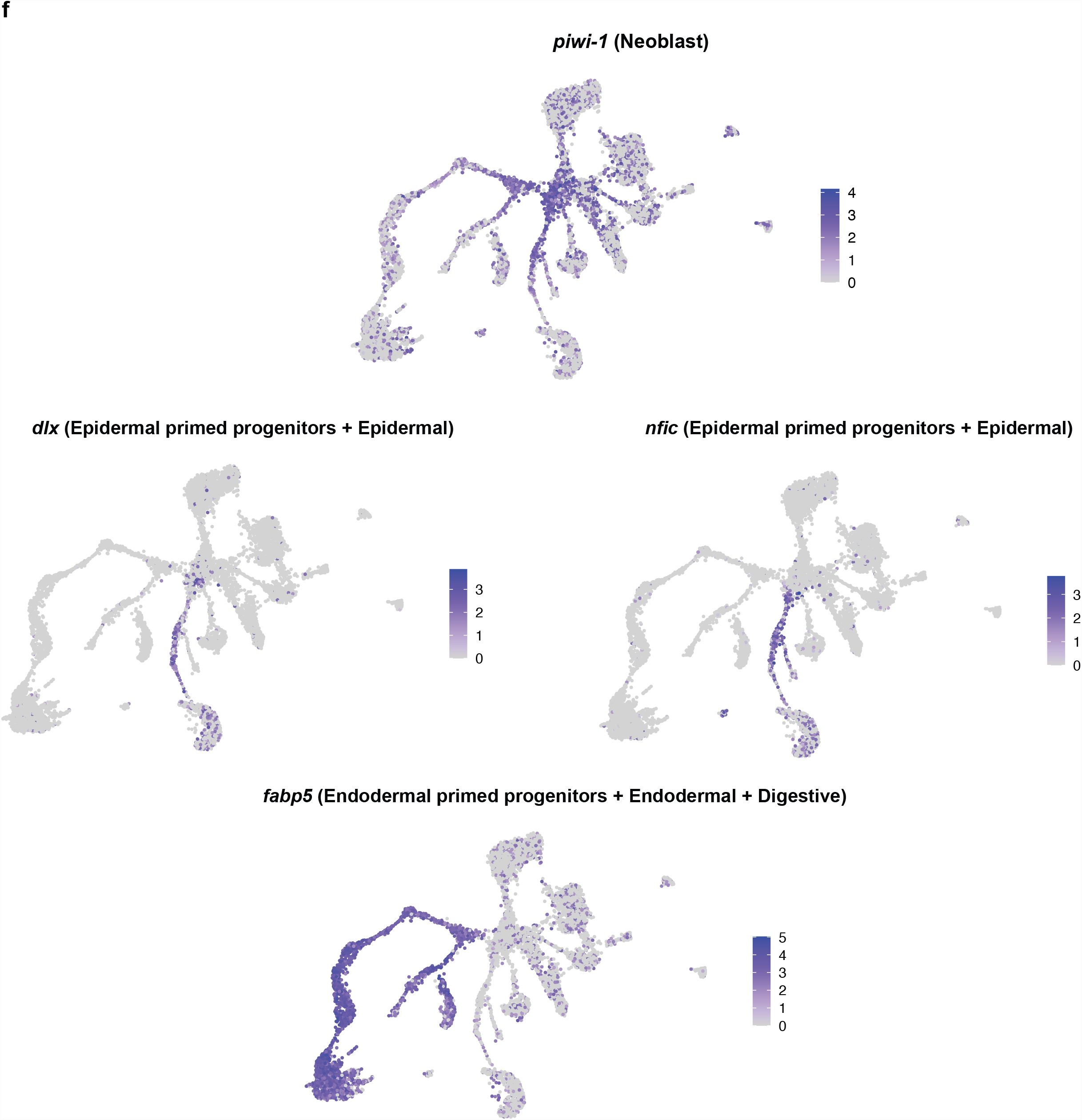
Identification and corroboration of shared cell types across postembryonic development. (A) UMAPs representing putative cell types across postembryonic development. Cluster markers for each data set can be found in Supplementary Table 1. (B) Heatmap of Pearson correlation coefficients based on transcriptional profiles of all cell clusters from the postembryonic data sets with hierarchical clustering. HJ= hatchling juvenile, LJ= late juvenile, EA= early adult, LA= late adult (C) UMAPs representing shared cell types across postembryonic development. Transcriptionally similar clusters are denoted by the same color across stages. (D) On the left: hatchling juvenile umap with putative cluster labels. On the right: Gene ontology (GO) terms associated with each hatchling juvenile cluster based on highly expressed genes found within each cluster. (E) FISH of genes associated with each cluster, with markers from the same cell clusters showing similar gene expression patterns. Scale bars 100 µm. Scale bars for germ line *in situs pa1b3-2* and *hspb1*: 300 µm and *ldlr6*: 500 µm. (F) UMAP showing *piwi-1* gene expression with enrichment in the central neoblast cluster as well as the putative epidermal and endodermal lineage-primed progenitors. UMAPs of Epidermal lineage-primed progenitor and differentiated populations depicted with *dlx* and *nfic* expression. UMAP of endodermal lineage-primed progenitor and differentiated endodermal + digestive cells depicted with *fabp5* expression.

**Supplementary Figure 2.**
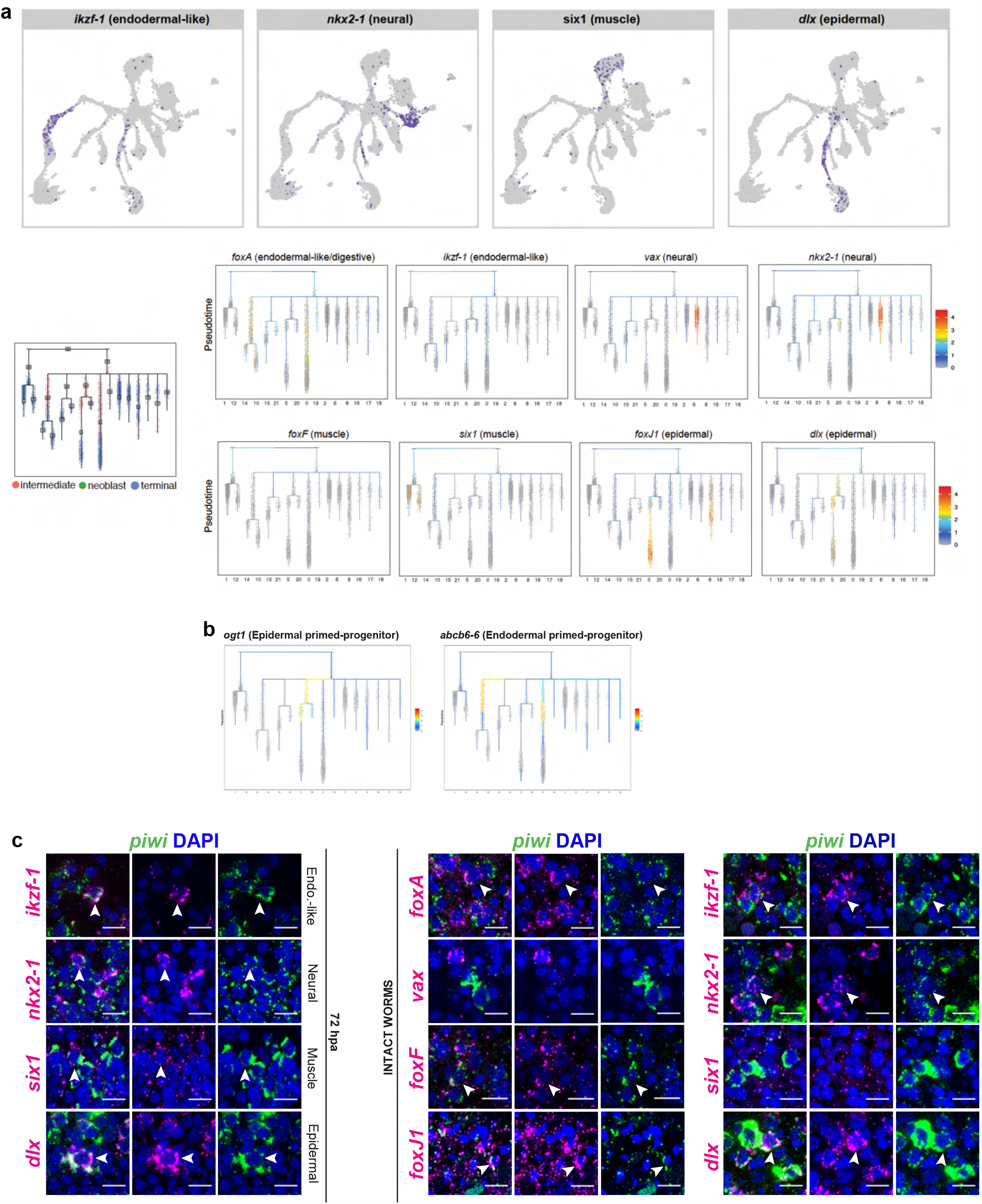
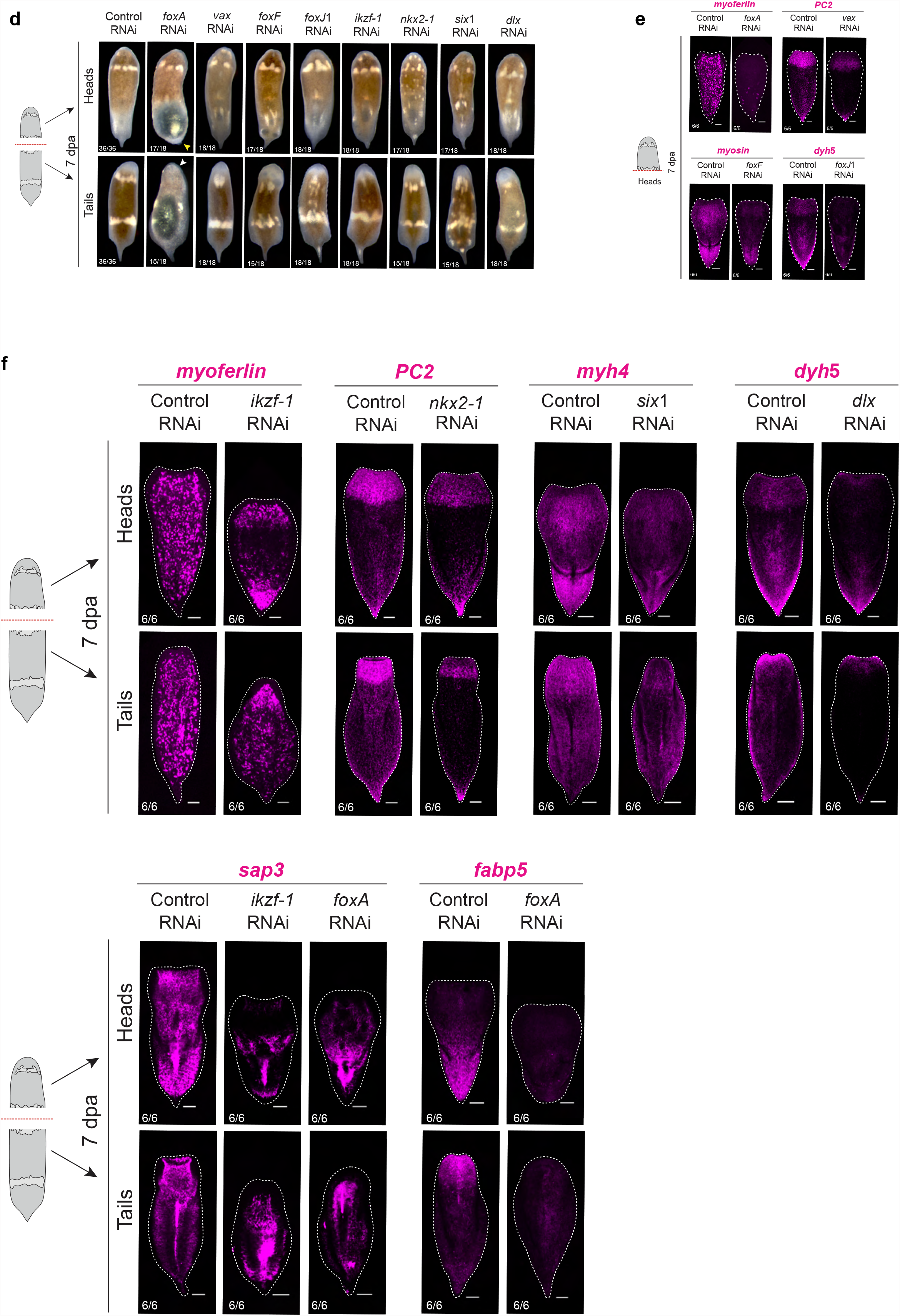
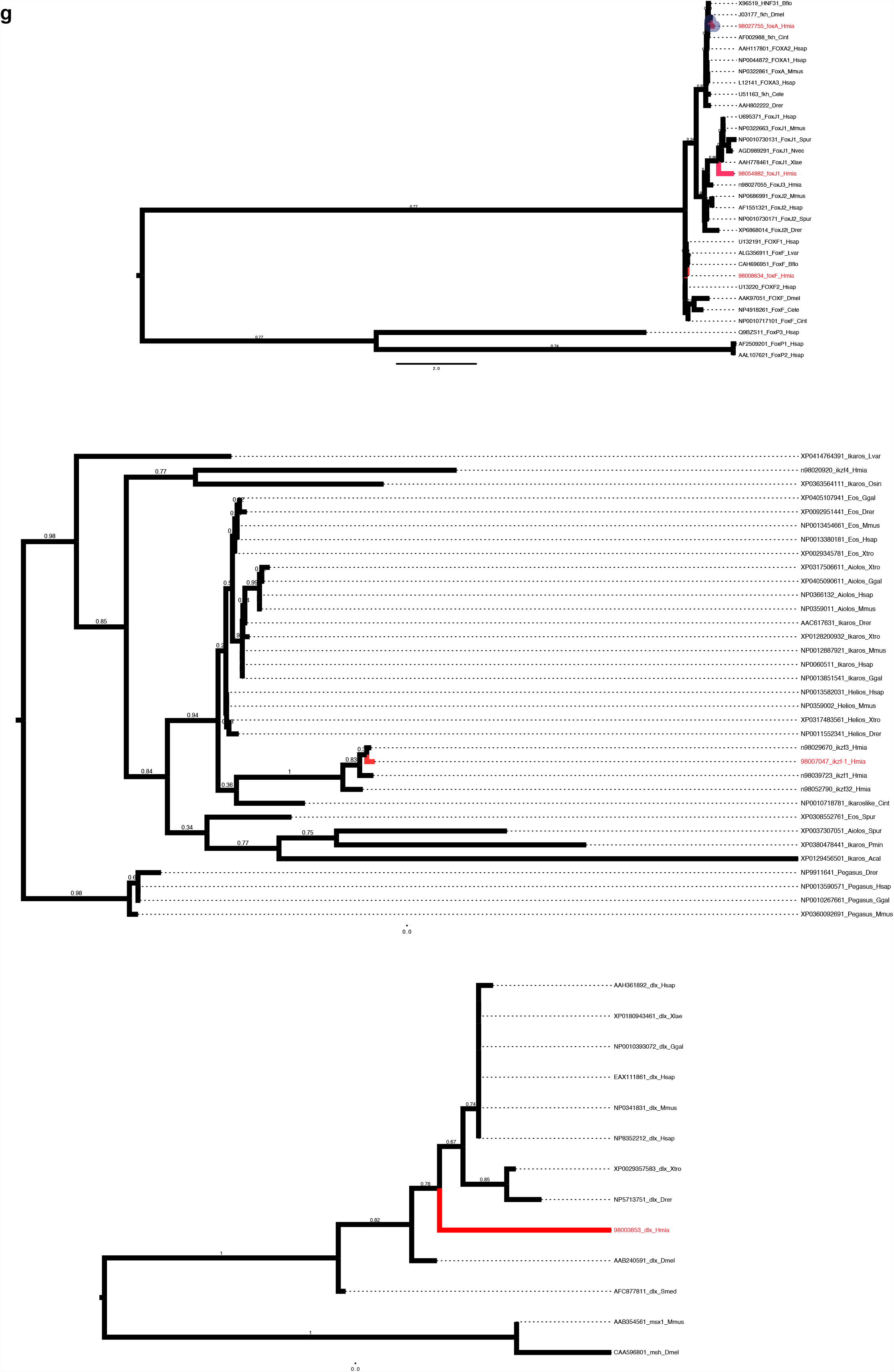
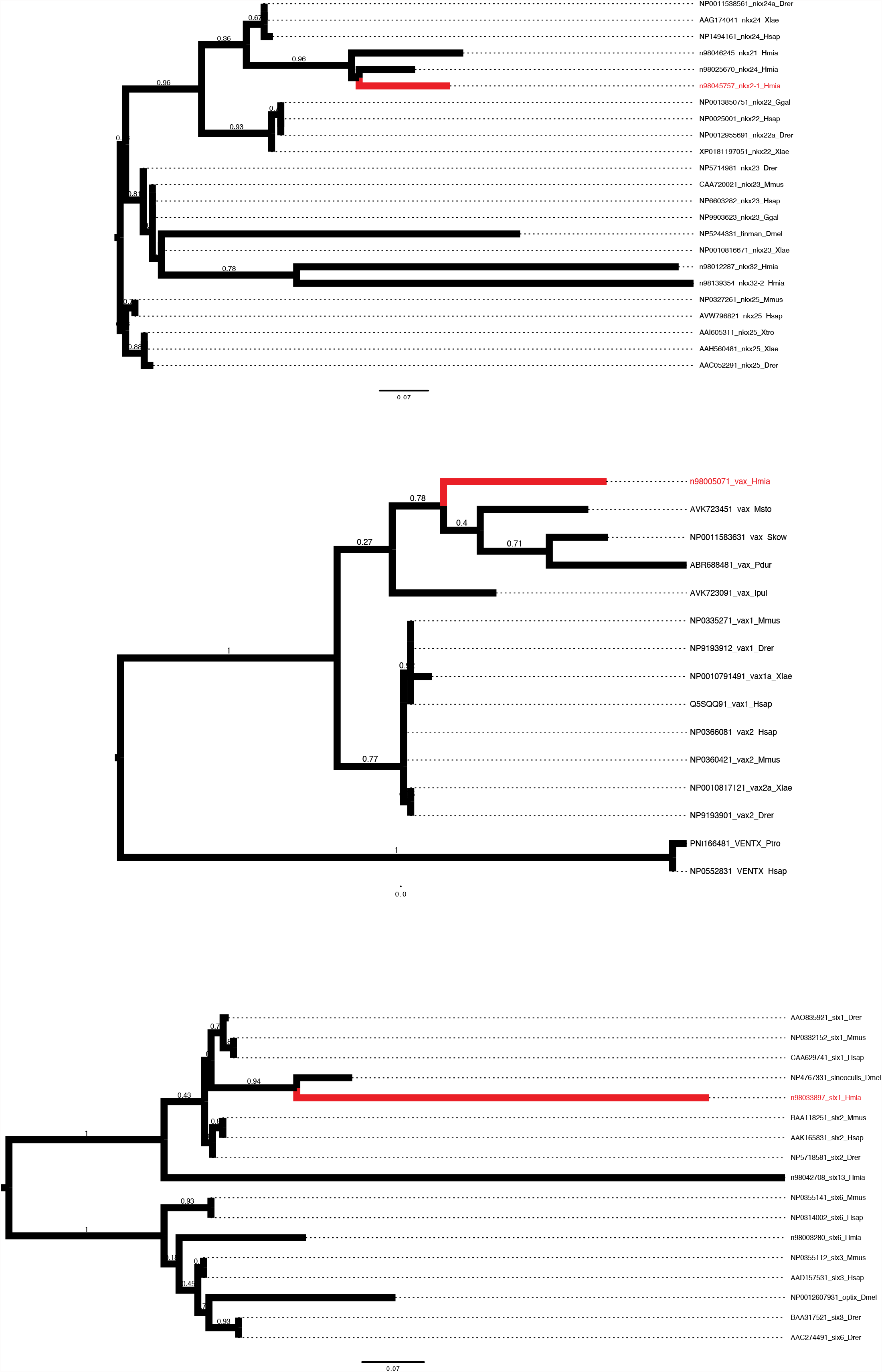
Additional transcription factors expressed in putative specialized neoblast populations with gene expression studies. (A) UMAP projection of transcription factors expressed in putative differentiation trajectories from URD analysis along with URD trajectories. URD schematic on the left depicts intermediate (pink), neoblast (green), and terminal (blue) cell assignments for trajectory inference of hatchling juvenile data. All transcription factors chosen as candidates were found to be expressed in their corresponding differentiated cell types and/or putative specialized neoblast populations. (B) URD trajectories of putative lineage-primed progenitors for the epidermal and endodermal lineages were placed in close relationship to differentiated epidermal and digestive cells. (C) Double FISH of the candidate transcription factors with the stem cell marker *piwi-1* assessing co-expression (denoted by white arrowheads) in intact animals and during regeneration at 72hpa putative endodermal-like/digestive, neural, muscle, and epidermal specialized neoblasts. Scale bars 10 µm. (D) External morphological assessment of regenerating fragments 7dpa shows animals with blastemas and regenerated heads and tails in every RNAi condition except for *foxA* RNAi, where animals failed to regenerate head blastemas (denoted by white arrowhead) and pointed tails (denoted by yellow arrowhead). (E) RNAi of putative specialized neoblast transcription factor leads to loss of differentiated cell type expression during regeneration. Regenerating head fragments are shown 7dpa corresponding to the tail fragments in figure 2c. Scale bars 100 µm. (F) Additional RNAi of putative specialized neoblast transcription factor leads to loss of differentiated cell type expression during regeneration. Regenerating head and tail fragments are shown 7 day post amputation (7dpa) with cell-type gene expression denoted in magenta. Scale bars 100 µm. (G) Phylogenetic trees for transcription factors associated with putative specialized neoblasts.

**Supplementary Figure 3.**
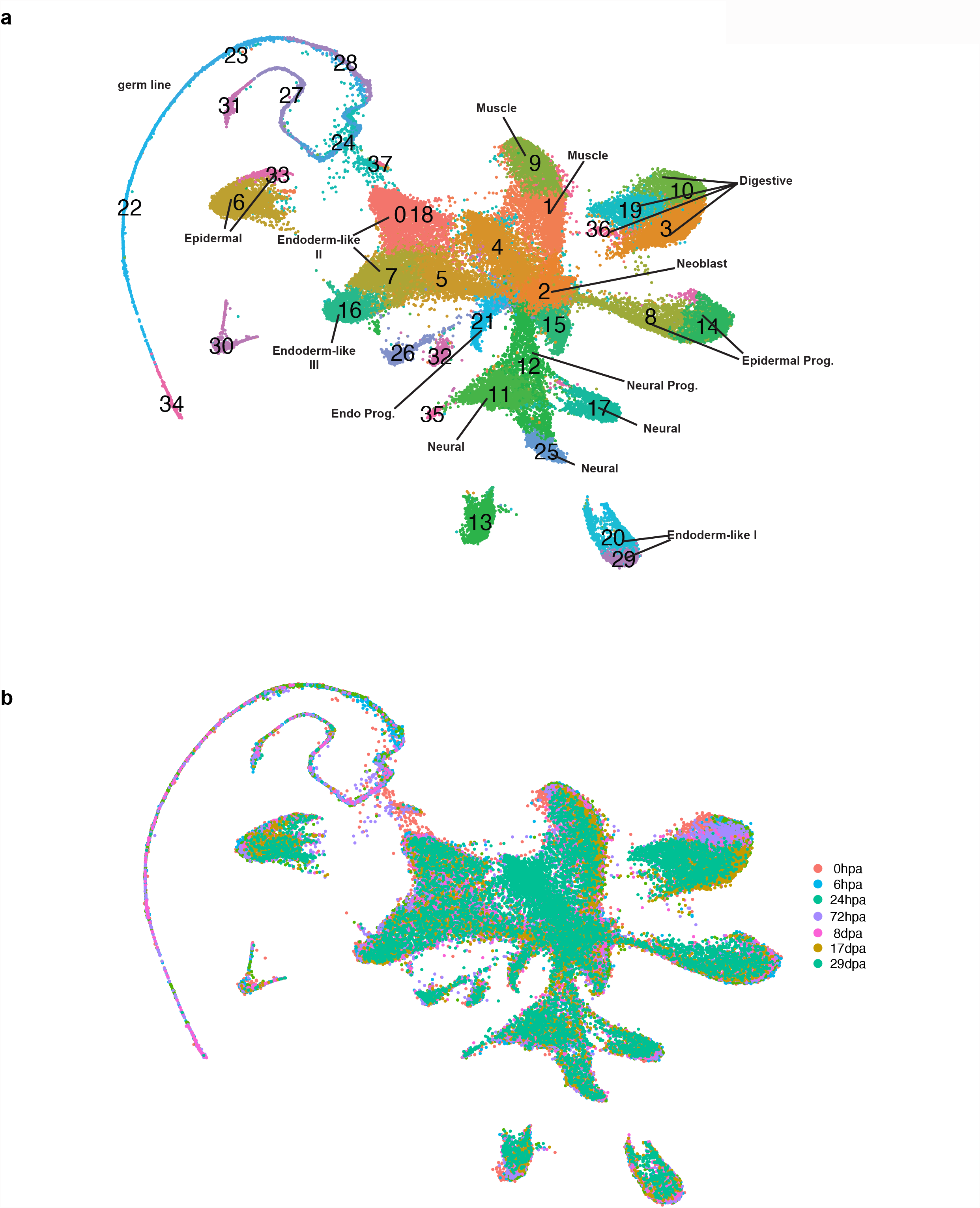
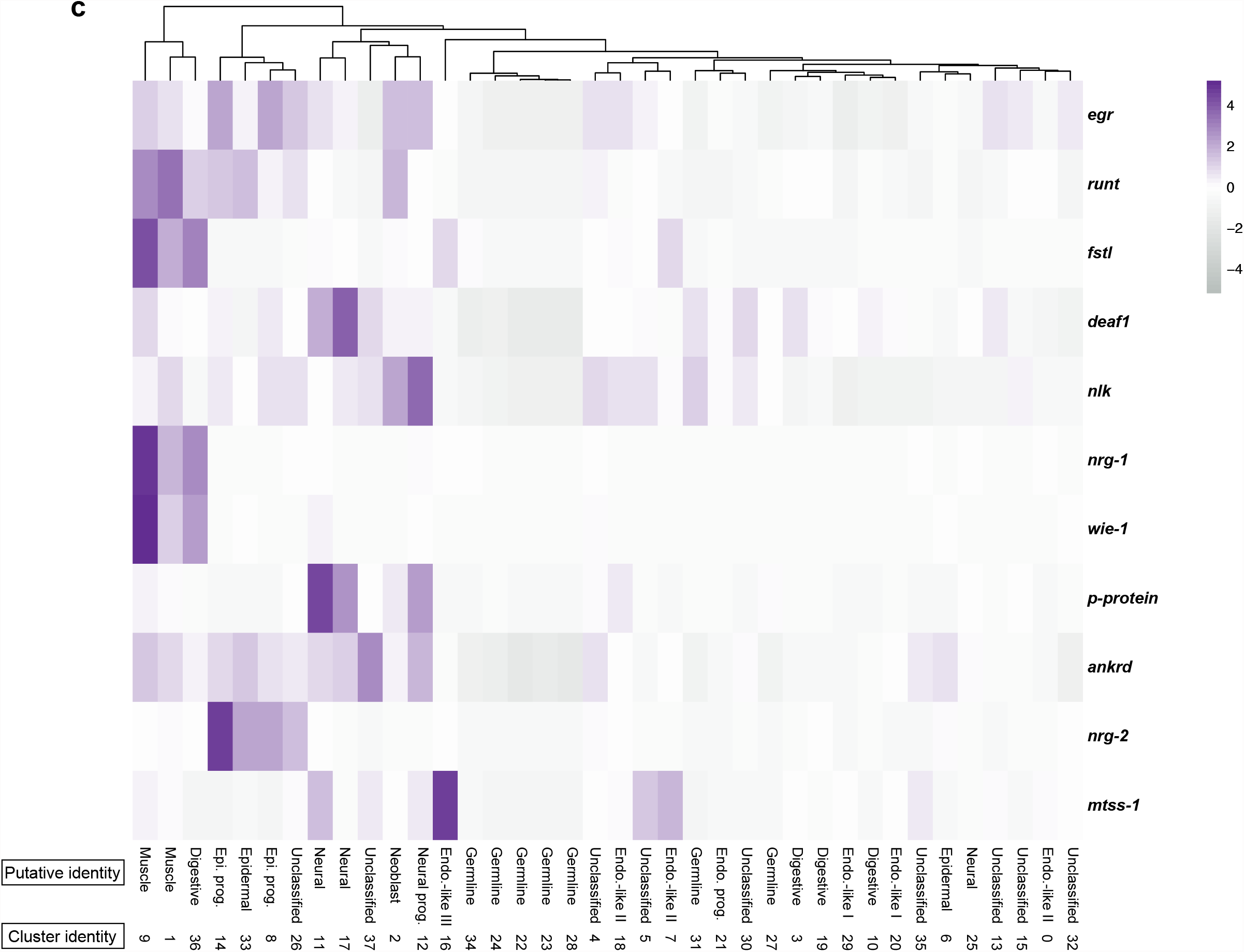
Integrated regeneration UMAP of scRNA-seq and cluster-specific responses to the Egr-GRN. (A) UMAP of integrated regeneration scRNA-seq data annotated with cell clusters identified through unsupervised clustering and shared cell types identified in the postembryonic data sets. (B) UMAP of integrated regeneration scRNA-seq data was composed of cells from each of the regeneration time points, with no regeneration-specific cluster clearly identifiable. (C) Cluster-specific expression of Egr-GRN. Cells from the integrated regeneration data were used to generate a heatmap depicting average expression, with putative identity labeled based on similarity to the shared cell types from the postembryonic data and cluster number from the UMAP in Supplementary Figure 3a.

**Supplementary Figure 4.**
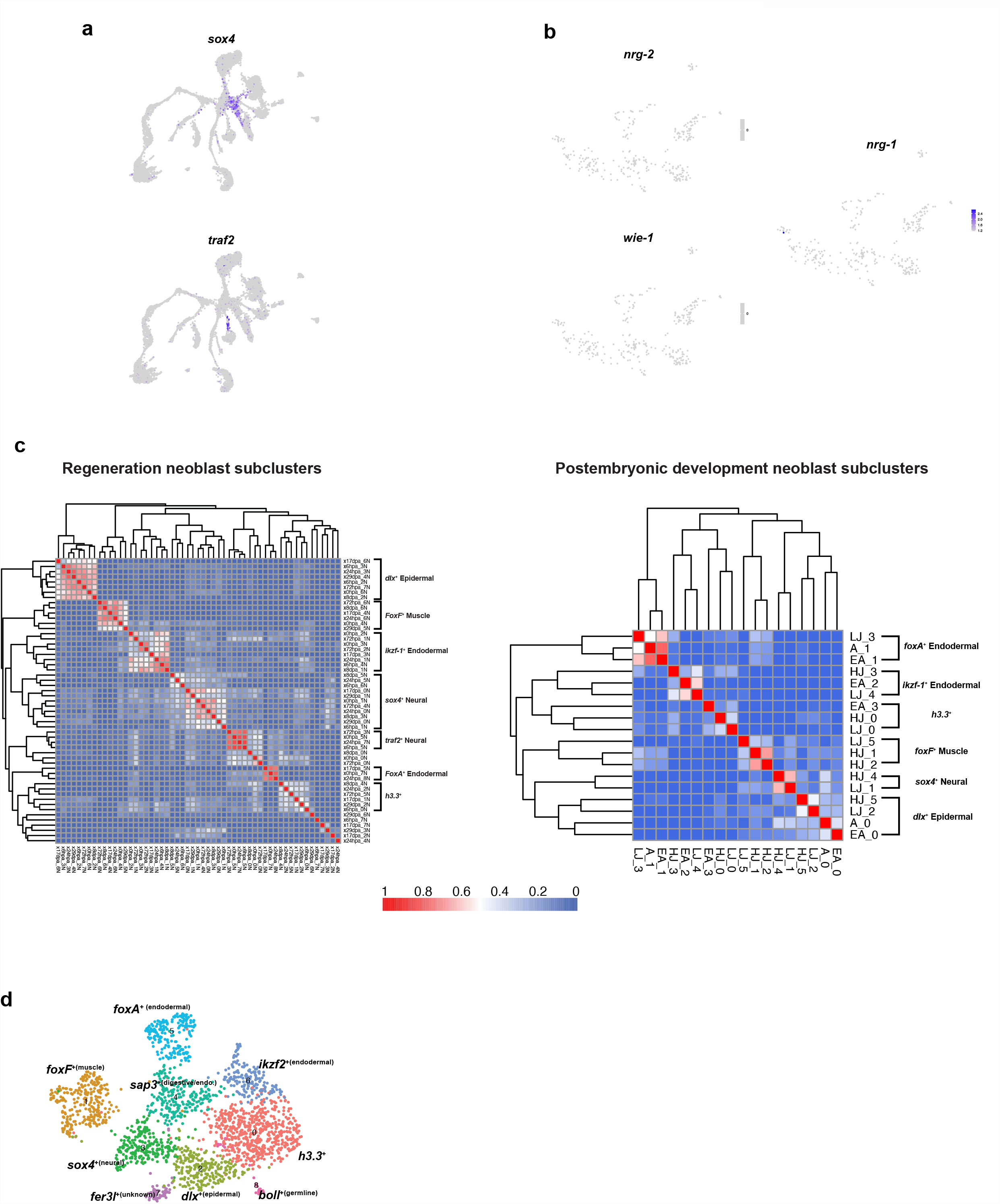
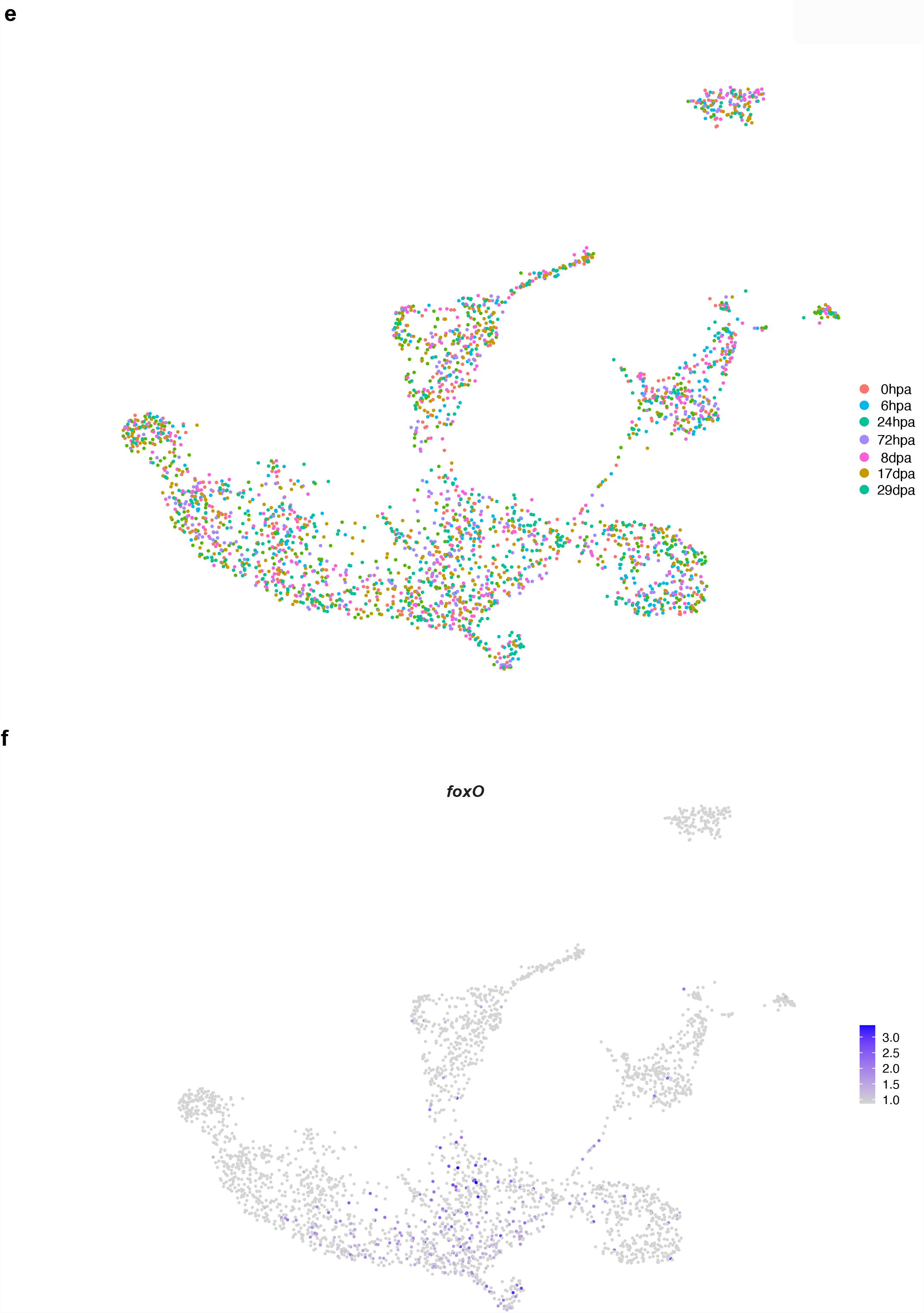
Specialized neoblast in postembryonic development and regeneration. (A) Projection of *sox4* and *traf2* genes in the hatchling juvenile UMAP supports their putative neural identity. (B) Egr-GRN members *nrg-2, nrg-1*, and *wie-1* are not expressed in neoblasts at 6hpa in Fig. 4b so they were not included in the heatmap. (C) Heatmaps of Pearson correlation coefficients based on transcriptional profiles of all neoblast subpopulations from the regeneration and postembryonic neoblasts. (D) UMAP of integrated postembryonic development neoblast subpopulations with differentially expressed marker gene overlaid. (E) UMAP of integrated regeneration neoblast subpopulations showing that clusters are composed of cells from each of the regeneration time points, including the *h3*.*3*^+^ population. (F) UMAP of integrated regeneration neoblast subpopulations showing *foxO* gene expression.

**Supplementary Table 1**. Top markers from each cluster for the postembryonic development stages.

**Supplementary Table 2**. Orthology assessment for transcription factors associated with specialized neoblast and progenitor populations.

**Supplementary Table 3**. Top markers from each cluster for the regeneration stages.

## Methods and Materials

### Methods

#### Animal maintenance

Gravid adult worms were kept in plastic boxes (20–30 worms per 2.13-L Ziploc container) at 21°C in artificial seawater (38 ppt, pH 7.9–8.0; Instant Ocean Sea Salt). Juvenile worms were kept in zebrafish tanks (∼300 worms per tank). Seawater was replaced twice a week. Juvenile and adult worms were fed with rotifers *Brachionus plicatilis* and freshly hatched brine shrimp *Artemia sp*. twice a week, respectively.

#### Cell dissociation and preparation

To avoid cell loss, juvenile and adult worms were dissociated in calcium- and magnesium-free artificial seawater (CMF-ASW: 450 mM NaCl, 9 mM KCl, 30 mM Na_2_SO_4_, 2.5 mM NaHCO_3_, 25 mM HEPES, 10 mM Tris-HCl, 2.5 mM EDTA) in a 2-mL DNA LoBind tube (Eppendorf) by vigorously pipetting using a P1000 micropipette. Cell suspensions were passed through a 40-μm cell strainer (Falcon) to remove remaining aggregates. To further remove cell debris and cell-free RNA, cells were loaded on a BSA cushion (4% BSA in CMF-ASW) and pelleted at 500 ×*g* for 5 min using a swing rotor. Cells were then washed twice with CMF-ASW by resuspension and spinning at 500 ×g for 3 min. For inDrops encapsulation, cell suspensions in CMF-ASW were mixed 1:1 with 30% OptiPrep in CMF-ASW.

#### inDrops encapsulation and library preparation and RNA sequencing

Encapsulation of single cell suspensions were performed at the Single Cell Core located at the Harvard Medical School using a custom inDrops microfluidics system (Zilionis et al., 2017). *Hofstenia* is a marine organism, meaning it was necessary to keep the cells in artificial seawater until the last possible moment before encapsulation to prevent cell death. We utilized a custom microfluidics chip offered at the core that allowed for the introduction of 1xPBS solution just prior to encapsulation. Library preparation was performed by the Single Cell Core as well. Library quality control via tapestation, qPCR quantification, and sequencing using the Illumina Nextseq (75 bp, paired-end) was performed by the Harvard Bauer Core.

#### Single-cell analysis

Once reads were acquired, they were demultiplexed, mapped, and subsequently converted into count matrices using the methods described in the following github repository: https://github.com/brianjohnhaas/indrops. Cell quality filtering, clustering, and dimensionality reduction was done using Seurat v3 in R(Butler et al., 2018; Hao et al., 2021; Satija et al., 2015; Stuart et al., 2019). All of the code used to analyze the different datasets will be available on our github repository.

#### Fixation and in situ hybridization

Whole worms and regenerating fragments were fixed in 4% paraformaldehyde in 1% phosphate-buffered saline with 0.1% Triton-X (PBST) for one hour at RT on nutator and stored in methanol at -20 °C until use. Digoxigenin and Fluorescein labeled riboprobes were synthesized as previously described (Srivastava et al., 2014). Fluorescence in situ hybridizations (FISH) were performed following the protocol described in (Srivastava et al., 2014) with minor modifications. Animals were washed twice for ten minutes each in PBST and then blocked in a 10% horse serum in PBST for one hour at RT before antibody incubation. All washes the following day were made with PBST followed by tyramide with rhodamine or fluorescein development for 10 min. Detailed probe synthesis and *in situ hybridization* protocols with reagents and solution preparations are found in (Srivastava et al., 2014).

#### RNAi

Double-stranded RNA (dsRNA) synthesis was made following a protocol previously described in (Srivastava et al., 2014). RNAi experiments were done by injecting the animals with dsRNA corresponding to the target gene into the gut for 3 consecutive days. Animals were cut transversally at least 2 hours after the third injection and were allowed to regenerate for 7 days while being monitored for visible phenotypes or external defects. Control dsRNA for gene inhibition was the *unc22* sequence from *C. elegans* that is absent in *Hofstenia*. After 7 days post-amputation (dpa) animals were fixed and analyzed by FISH. dsRNA injections were performed using a Drummond Nanoject II.

#### Gene ontology enrichment analysis

Gene ontology enrichment analysis was done using the same methods described in (Kimura et al., 2021).

#### Lineage reconstruction

Lineage reconstruction was done using the R package URD(Farrell et al., 2018). To construct lineage trees at the Hatchling Juvenile stage, we set the neoblast cluster as the root, differentiated cell types (i.e. digestive, epidermal, neural, muscle) as the lineage tree tips, and all other cells as the intermediate cells. All of the parameters utilized for constructing the URD tree along with differential expression will be available on our github repository.

#### Pseudo-bulk heatmap

Cellular populations were subsetted from integrated UMAPs based on clustering (i.e. the muscle cluster was extracted and further analyzed). Metadata associated with each data set, including post embryonic developmental stage and regeneration time point, were used to subset and further characterize expression dynamics.

#### Phylogenetic analysis

Transcription factors identified from the scRNA-seq data, URD trajectory inference, and whose function were subsequently tested were assigned orthology. First, a BLAST was performed to putatively identify transcription factors of interest. To assign orthology, phylogenetic trees were constructed. Sequences were aligned with MUSCLE (v3.8.31) (Edgar, 2004). Alignments were trimmed using Gblocks (Castresana, 2000; Talavera & Castresana, 2007) with the least stringent parameters Phylogenetic trees were inferred using Maximum Likelihood analysis with 1,000 bootstrap replicates, implemented in RAxML (v8.2.4) (Stamatakis, 2014) using the WAG+G model of protein evolution.

## Acknowledgements

We want to thank the Harvard Medical School Single Cell Core for encapsulation and library preparation. As well as the Harvard Bauer Core for their assistance in sequencing the libraries. D. Wagner, A. Veres, and J.A. Farrell for help and guidance in data analysis. Thank you to members of the Srivastava and Koenig labs for thoughtful discussion and scientific insight. V. Beinhart for corroboration of clustering between Seurat v2 and v3. V. Chandra, K. Loubet-Senear, C. Breen, and A.Q. Rock for critical reading of the manuscript.

## Competing interests

No competing interests declared.

## Funding

This work was supported by grants to M.S. by the Searle Scholars Program, Smith Family Foundation, and the National Institutes of Health (1R35GM128817-01). R.E.H. is supported by the National Institutes of Health F31 (1F31GM134633-01A1). J.O.K. is supported by the NSF-Simons Center for Mathematical and Statistical Analysis of Biology at Harvard and the Harvard Quantitative Biology Initiative (1764269). Y-J.L is supported with the HFSP Long-Term Fellowship (LT000022/2017-L)

## Author Contributions

Y-J.L., R.E.H, L.R., and M.S. conceived the study. R.E.H., J.O.K., D.M.B., and M.S. wrote the paper. Y-J.L. generated sequence data, data matrices, and conducted preliminary analysis. J.O.K. and R.E.H. performed subsequent single-cell data analysis and clustering. D.M.B., R.E.H., and L.R. performed FISH corroboration. D.M.B. and R.E.H. performed confocal imaging. J.O.K. performed URD lineage reconstruction. D.M.B. performed RNAi of transcription factors of interest. R.E.H. performed phylogenetic analysis of transcription factors of interest.

## Data availability

Sequencing raw reads and processed counts matrices associated with this study will be deposited in the NCBI Gene Expression Omnibus (GEO). An online resource generated to access and visualize the scRNA-seq data will be provided.

## Notes

### Competing Interest Statement

The authors have declared no competing interest.

